# MLF2 modulates phase separated nuclear envelope condensates that provoke dual proteotoxicity

**DOI:** 10.1101/2021.10.26.465916

**Authors:** Sarah M Prophet, Anthony J Rampello, Robert F Niescier, Juliana E Shaw, Anthony J Koleske, Christian Schlieker

## Abstract

DYT1 dystonia is a highly debilitating neurological movement disorder arising from mutation in the AAA+ ATPase TorsinA. The hallmark of Torsin dysfunction is nuclear envelope blebbing resulting from defects in nuclear pore complex biogenesis. Whether blebs actively contribute to disease manifestation is presently unknown. We report that FG-nucleoporins in the bleb lumen undergo phase separation and contribute to DYT1 dystonia by provoking two proteotoxic insults. Short-lived ubiquitinated proteins that are normally rapidly degraded in healthy cells partition into the bleb lumen and become stabilized. Additionally, blebs selectively sequester a chaperone network composed of HSP70s and HSP40s. The composition of this chaperone network is altered by the bleb component MLF2. We further demonstrate that MLF2 is a catalyst of phase separation that suppresses the ectopic accumulation of FG-nucleoporins and modulates the selective properties and size of condensates *in vitro*. Our studies identify unprecedented, dual mechanisms of proteotoxicity in the context of liquid-liquid phase separation with direct implications for our understanding of disease etiology and treatment.

## Introduction

Torsin ATPases (Torsins) are the only members of the AAA+ protein superfamily that localize within the endoplasmic reticulum (ER) and nuclear envelope (NE)^1, 2^. TorsinA is essential for viability^3^ and strictly requires regulatory cofactors to hydrolyze ATP^4^. A mutation in TorsinA that disrupts the interactions with its cofactors results in a loss of ATPase activity ^5, 6^ and is responsible for a debilitating neurological movement disorder called DYT1 dystonia ^7^. The Torsin activator LAP1 was recently implicated in NE dynamics^8^, and LAP1 mutations give rise to dystonia and myopathy^9–11^. Thus, the Torsin system is critical for NE dynamics and neurological function^10, 12^. While the molecular target of Torsin activity and the mechanism of DYT1 dystonia onset remain to be identified, the ubiquitous phenotype observed in diverse DYT1 dystonia models is NE blebbing^3, 13–22^. Our group reported that NE blebs result from defective nuclear pore complex (NPC) biogenesis^23^ and consistently, nuclear transport defects have been observed both in organisms and cells with compromised Torsin function including patient- derived iPSC neurons^13, 21, 22^. NPCs are composed of proteins called nucleoporins (Nups), several of which contain disordered phenylalanine-glycine (FG)-rich domains that establish the permeability barrier characteristic of NPCs^24–26^. The native disorder and amino acid composition of FG-Nups allows them to phase separate within the NPC’s central channel through which small (<30kDa) molecules can passively diffuse^27^. Larger molecules require facilitated passage via nuclear transport receptors (NTRs)^28^.

NE blebs arising from Torsin deficiency are enriched for FG-Nups, but do not contain NTRs or bulk nuclear export cargo^19, 23^. Moreover, the poorly characterized myeloid leukemia factor 2 (MLF2) protein and K48-linked ubiquitin (Ub) chains are diagnostic constituents of the bleb lumen^19, 21, 23^. While Ub accumulation and defects in the ubiquitin/proteasome system have been implicated in many other neurological disorders including Huntington’s and Parkinson’s disease^29–31^, it is generally unknown whether or how NE blebs interfere with cellular protein quality control (PQC). Both the lack of suitable readouts and our incomplete understanding of the molecular composition of NE blebs represent major obstacles towards identifying functional consequences of the interplay between FG-Nups, MLF2, and the PQC system.

In this study, we develop a viral model substrate to define the bleb proteome and to probe the hitherto unknown significance of ubiquitin accumulation for disease etiology. We find that normally short-lived proteins evade degradation once they are trapped inside blebs. Along with these stabilized proteins, blebs selectively sequester a specific chaperone network composed of HSP40s and HSP70s both in human cell lines and in primary murine neurons. The formation of blebs requires the FG-Nup Nup98 to form a phase-separated compartment within the bleb lumen. Furthermore, we combine cellular and *in vitro* approaches to assign a role to MLF2 as a modulator of FG-Nup phase separation. MLF2 catalyzes the formation of large condensates and modulates their permissiveness towards NTR-like molecules. Together, our results advance our understanding of cellular phase separation and define an unprecedented link between PQC defects and disease etiology via a pathological, phase separated NE compartment.

## Results

### Torsin deficiency causes the stabilization of proteins that are rapidly degraded in WT cells

To develop approaches for exploring the poorly understood consequences of protein sequestration into NE blebs, we examined viral proteins that have been functionally tied to nuclear transport. ORF10 from Kaposi’s sarcoma-associated herpesvirus (KSHV) is produced as a full length 418-residue protein and, as often observed in viral proteins, as a shorter 286-133 ORF10 via alternative translation initiation (Fig. 1a,b). The full-length protein inhibits mRNA export by binding to the export factor Rae1 during the KSHV lifecycle^32^ and localizes diffusely within the nucleoplasm in wild type (WT) and TorsinKO HeLa Δ133 ORF10 becomes tightly sequestered into NE foci that strictly co-localize with K48-Ub in TorsinKO cells (Fig. 1c). Because nucleoplasmic in WT cells (Fig. 1c) and is associated with more K48-Ub in TorsinKO compared to WT cells (Fig. 1d), we conclude that ubiquitylated This recruitment depends on ubiquitylation: fusing the virus-derived M48 deubiquitylating (DUB) activity^33^ to Δ133 ORF10 prevents NE sequestration, while a catalytically inactive DUB variant has no effect (Fig. 1g).

**Figure 1.**
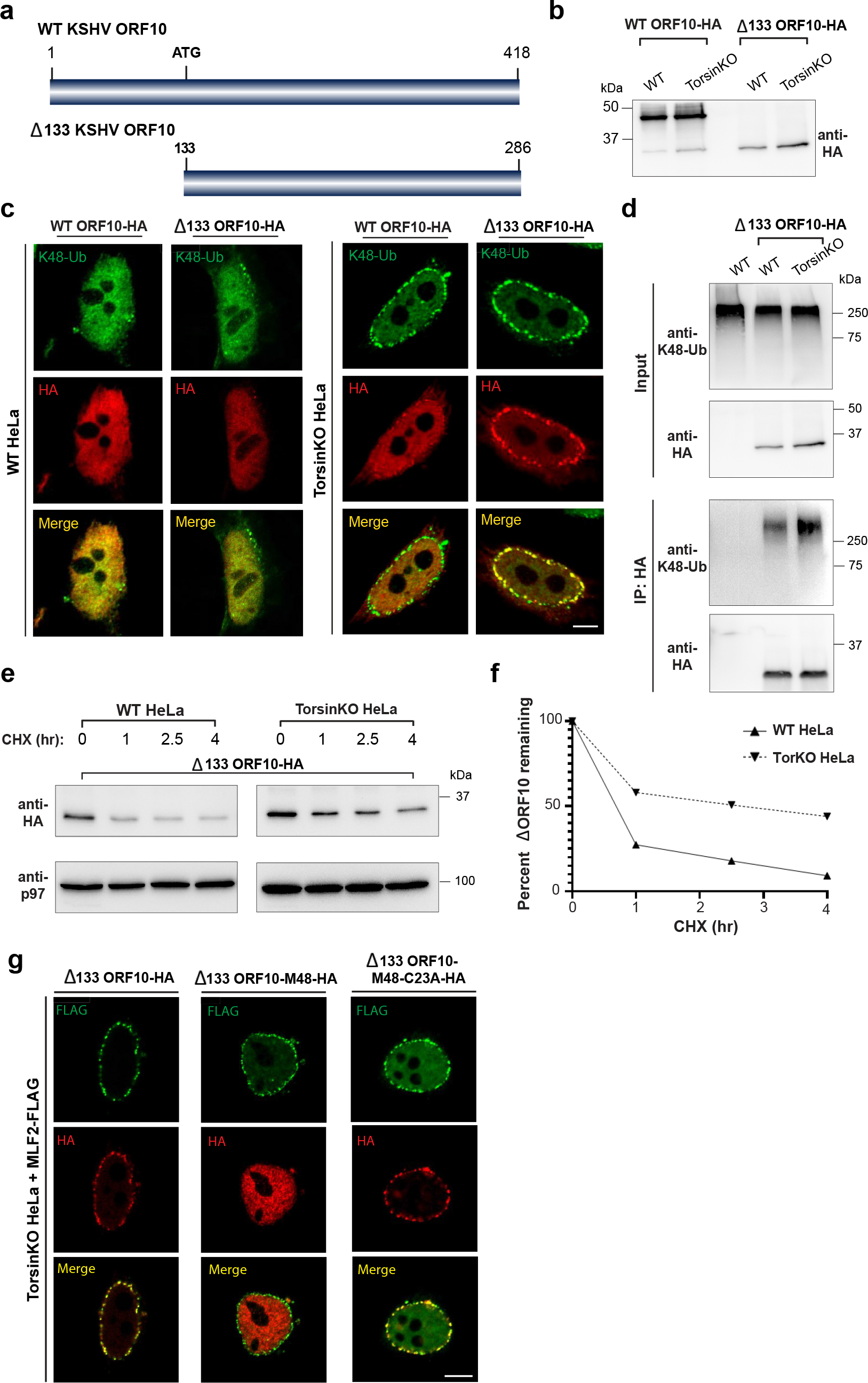
NE herniations arising from Torsin ATPase deficiency sequester and stabilize short-lived protein. **a**, Schematic model of the ORF10 protein from KSHV. KSHV ORF10 contains an internal start codon at residue 133 that produces Δ133 ORF10. **b**, Immunoblot using anti- HA demonstrating expression of Δ133 ORF10-HA in WT and TorsinKO HeLa cells 24 hours post transfection. Note that ORF10-HA is produced as a major full-length protein a lower abundance Δ133 product. **c**, Representative IF images of full length and Δ133 ORF10-HA in WT and TorsinKO cells. Scale bar, 5 µm. **d**, Anti-HA IP from WT or TorsinKO cells expressing Δ133 ORF10-HA. The IP was probed with antibodies against K48-Ub and HA. Note that Δ133 ORF10-HA is associated with more K48-Ub in TorsinKO than WT cells. **e**, A cycloheximide (CHX) chase over four hours in WT and TorsinKO cells expressing Δ133 ORF10-HA. Cells were treated with 100 µg/mL of CHX at 37°C for the indicated timepoints. p97 serves as a loading control. **f**, Relative percentage of Δ133 ORF10-HA obtained in (**e**) was determined via densitometry by comparing to the abundance at time = 0. All data were standardized to p97 levels. **g**, Representative IF images of TorsinKO cells expressing MLF2-FLAG and Δ133 ORF10-HA fusion constructs. Δ133 ORF10 was fused to the cytomegalovirus deubiquitinase (DUB) domain, M48 (ref.59). M48 is a highly active DUB domain that efficiently removes ubiquitin conjugates (ref.23, 33). A C23A mutation renders the DUB domain catalytically inactive (ref.59). Scale bar, 5 µm.

As Δ133 ORF10 is less abundant and associated with lower levels of K48-Ub in WT cells compared to TorsinKO cells (Fig. 1d), we hypothesized that short-lived protein. Indeed, we observed its half-life in WT cells to be approximately 45 minutes Δ133 ORF10 is stabilized and exists with a half-life of at least four hours (Fig. 1e,f). In conclusion, a protein that is normally destined for rapid degradation becomes strongly stabilized in Torsin-deficient cells after its Ub-dependent recruitment to the bleb. This reveals an unexpected proteotoxic property NE blebs and establishes Δ133 ORF10 as an ideal model substrate to explore the connection of NE blebs to disease etiology.

### Blebs are enriched for a specific chaperone network

The absence of a comprehensive, bleb-specific proteome is a major limitation in understanding the molecular underpinnings of NE bleb formation. We fused the engineered ascorbate peroxidase APEX2^34, 35^ to MLF2-HA and performed a biotin-based proximity labeling reaction (Fig. 2a). To control the APEX2-MLF2-HA protein level, we placed its expression under a doxycycline (DOX)-inducible promoter (Fig. 2b). The presence of biotin conjugates within blebs after the APEX reaction was verified by immunofluorescence (IF) (Fig. 2c) and the biotin-conjugating activity of APEX2 was confirmed via immunoblotting (Fig. 2d). After performing the APEX reaction, NE fractions were isolated from WT and TorsinKO cells. Biotinylated proteins were enriched via streptavidin-coated beads and identified by liquid chromatography-tandem mass spectrometry (LC-MS/MS) (Fig. 2e, Table S1). In parallel, we leveraged our discovery of Δ133 ORF10-HA followed by LC-MS/MS (Fig. 2e, Table S1). This allowed for a direct comparison of the bleb proteome obtained from two independent approaches with our previously published dataset of bulk nuclear envelope preparations after immunoprecipitation with a K48-Ub-specific affinity resin^23^. From these three datasets, we only considered proteins with an ≥ TorsinKO samples compared to WT (Fig. 2e).

**Figure 2.**
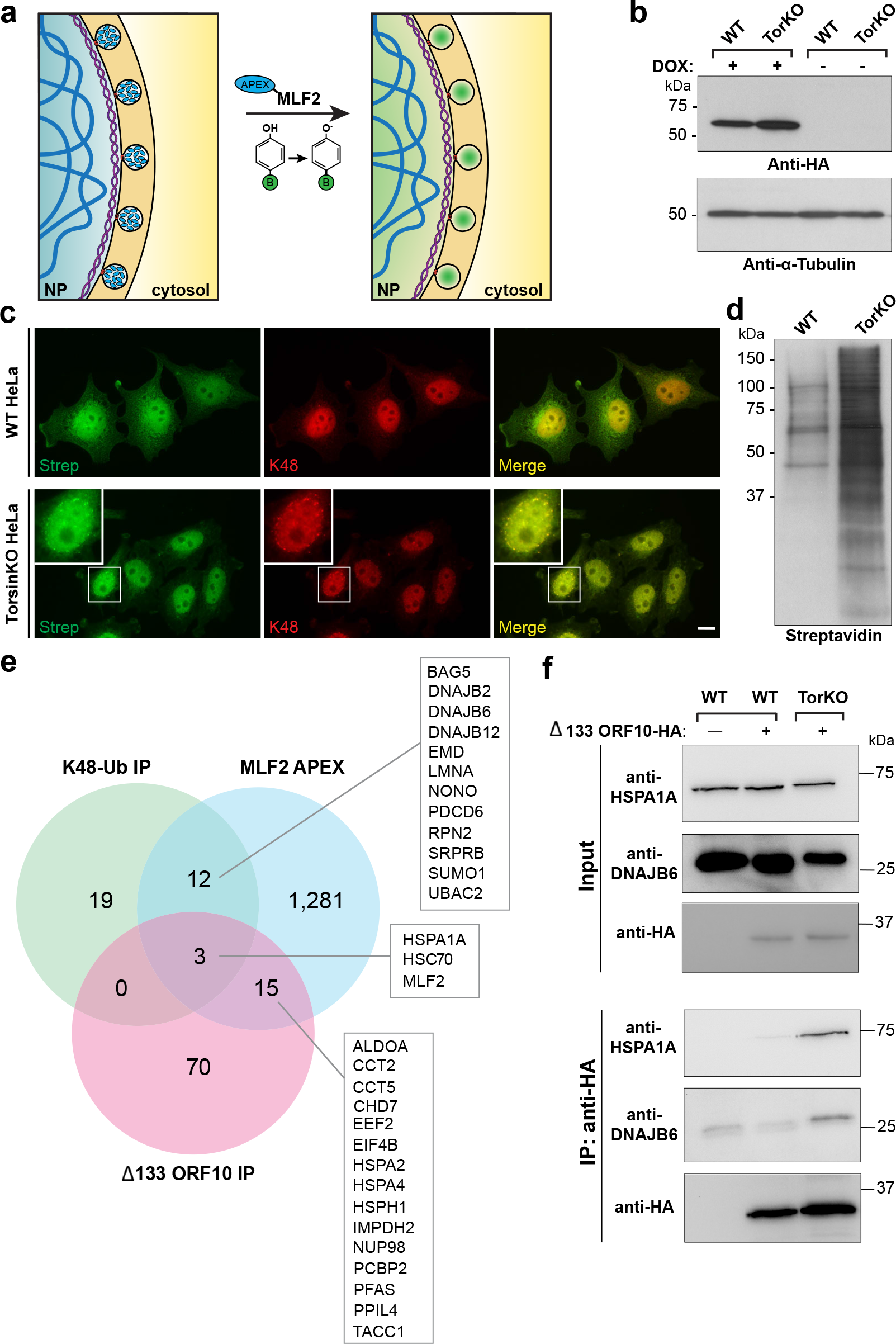
A comparative proteomics approach reveals NE blebs in Torsin-deficient cells are enriched for a highly specific chaperone network. **a**, A schematic illustration of the APEX2 reaction strategy to identify bleb protein contents. Left panel, the MLF2-APEX2 fusion protein (blue) localizes within the bleb lumen. Right panel, after incubation with 500 µM of biotin-phenol, cells are treated with 1 mM H2O2 and APEX2 oxidizes biotin phenol to form highly reactive biotin radicals that covalently label protein within a ∼20 nm radius (ref.60) (green cloud). NP, nucleoplasm. **b**, The expression of APEX-MLF2 was engineered in WT and TorsinKO cells to be under doxycycline (dox) induction. Cells were treated with dox for 24 hours before analysis by immunoblotting. **c**, Representative IF images of WT and TorsinKO cells after the APEX reaction. Note the enrichment of biotin conjugates in blebs of TorsinKO cells compared to the diffuse nuclear signal in WT cells. Strep (green) indicates fluorescently conjugated streptavidin signal and K48- Ub (red) indicate NE blebs. Scale bar, 10 µm. **d**, Immunoblot of NE fractions from WT and TorsinKO cells after the MLF2-APEX2 reaction as described in (**c**). **e**, Candidates were defined as proteins with spectral counts ≥1.5-fold enriched in TorsinKO compared to WT samples. The number of candidates identified for each of the three MS datasets are displayed as numbers within the Venn diagram. Hits overlapping between datasets are listed in alphabetical order. **f**, The stable interaction betweenΔ133 ORF10-HA and HSPA1A or DNAJB6 is unique to TorsinKO cells as judged by co- immunoprecipitation, consistent with the findings by comparative MS.

Only three proteins were consistently enriched across all datasets in samples from TorsinKO cells—MLF2, HSPA1A, and HSC70 (Fig. 2e). HSPA1A and HSC70 are the canonical cytosolic HSP70 members in mammalian cells, where they mediate a wide range of essential processes^36^. This functional diversity is achieved, at least in part, by interactions with J-domain proteins (HSP40s)^37^. Thus, we were interested to know whether the J-domain proteins shared between the APEX2-MLF2 and K48-Ub datasets may be enriched in blebs. First, we validated that Δ133 ORF10 stably interacts with DNAJB6 exclusively in TorsinKO cells by co-immunoprecipitation (IP) (Fig. 2f). These data suggest that beyond K48-Ub, MLF2, and FG-Nups, blebs harbor specific members of the HSP70 and HSP40 families.

### Members of the HSP70 and HSP40 families are sequestered into NE blebs

To validate our findings, we first tested whether overexpressed DNAJB12 and DNAJB2 localize to blebs by IF (Extended data 1a,b). While DNAJB12-HA showed relatively little recruitment to blebs (Extended data Fig. 1a), DNAJB2-HA strongly co-localized with K48-Ub NE foci (Extended data Fig. 1b) as we have previously observed^23^. Next, we determined whether HSPA1A, HSC70, DNAJB6, and DNAJB2 localized to blebs at endogenous expression levels using specific antibodies. In TorsinKO cells, HSPA1A, HSC70, DNAJB6, and DNAJB2 redistribute from diffuse cytosolic/nucleoplasmic distributions observed in WT cells (Fig. 3) to foci that decorate the nuclear rim (Extended data Fig. 1c, Fig. 3a). By overexpressing a dominant-negative TorsinA- EQ construct, we also determined whether these chaperones redistribute into blebs in the human neuroblastoma cell line SH-SY5Y (Fig. 3b). Indeed, upon TorsinA-EQ expression, HSPA1A and DNAJB6 became tightly sequestered into NE foci (Fig. 3b).

**Figure 3.**
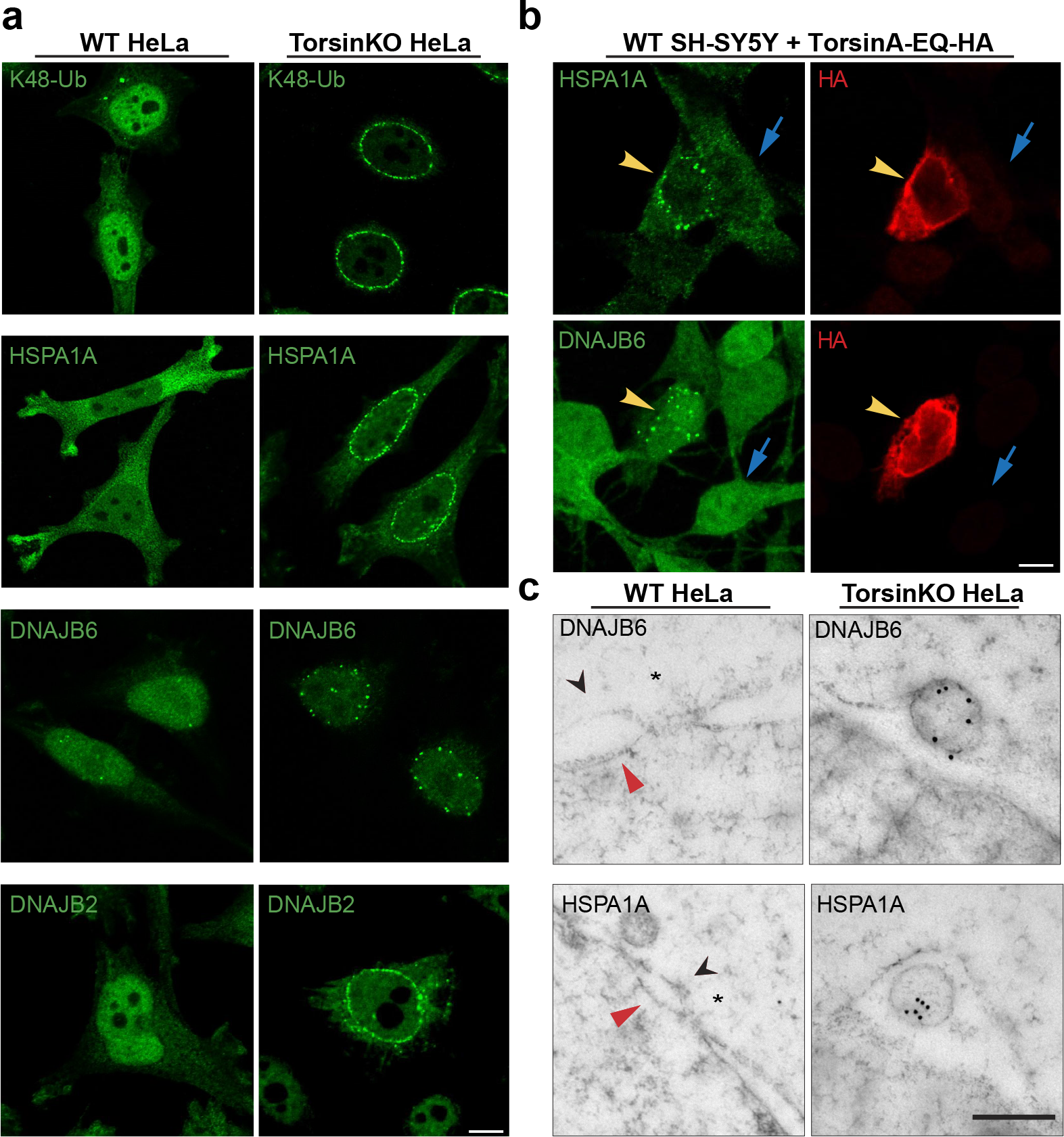
Highly abundant molecular chaperones are sequestered into NE blebs of Torsin-deficient tissue culture cells. **a**, Endogenous antibodies against K48-Ub, HSPA1A, DNAJB6, and DNAJB2 (green) reveal that chaperones from the HSP70 and HSP40 families become tightly sequestered into NE blebs upon Torsin deficiency. Scale bar, 5 µm. **b**, SH-SY5Y cells expressing a dominant-negative TorsinA construct, TorsinA- EQ-HA, sequester HSPA1A and DNAJB6 into NE blebs. Yellow arrowhead, transfected cell. Blue arrow, untransfected cell. Endogenous chaperones (green) form foci around the nuclear rim upon TorsinA-EQ-HA (red) expression. Scale bar, 5 µm. **c**, EM ultrastructure of the NE from WT or TorsinKO cells labeled with immunogold beads conjugated to anti-DNAJB6 (top) or anti-HSPA1A (bottom). Black arrowhead, outer nuclear membrane. Red arrowhead, inner nuclear membrane. Asterisk, NPC. Scale bar, 250 nm.

To confirm that these chaperones localize within the lumen NE herniations, we performed immunogold labeling and examined the ultrastructure of the NE with electron microscopy (EM) (Fig. 3c). We did not detect DNAJB6 or HSPA1A at mature NPCs in WT HeLa cells (Fig. 3c). However, in TorsinKO cells, DNAJB6 and HSPA1A localize within the bleb lumen (Fig. 3c). We conclude that multiple members of the HSP70 and HSP40 families become tightly sequestered into the bleb lumen in Torsin-deficient cells.

### Primary neurons with compromised TorsinA function sequester HSP70s and HSP40s into blebs

As DYT1 dystonia is a neurological disease, we were interested whether the sequestration of chaperones occurs in neurons with compromised Torsin function. In mouse models of DYT1 dystonia, ≥80% of central nervous system (CNS) nuclei exhibit NE blebs in eight-day-old mice^17^. This number drastically decreases as expression of TorsinB begins after about 14 days^17^. We therefore cultured mouse primary hippocampal neurons and introduced GFP with or without the dominant-negative TorsinA-EQ construct into primary hippocampal cultures after four days *in vitro* (DIV4). Cells were processed for IF on DIV7 (Fig. 4) to recapitulate the peak blebbing phenotype reported in conditional TorsinKO mice^17^. GFP was used to discern neurons from other cell types in our primary cultures based on cellular morphology. In neurons with functional Torsins, the chaperones are diffuse throughout the cytosol/nucleoplasm (Fig. 4a-d). Upon expression of TorsinA-EQ, these chaperones become sequestered into blebs (Fig. 4a-d). MLF2-HA is also sequestered into blebs in Torsin-deficient neurons even when overexpressed (Extended data Fig. 1d). Thus, the sequestration of highly abundant and essential molecular chaperones into blebs is a conserved and general consequence of Torsin dysfunction, suggesting a major role of blebs during disease manifestation.

**Figure 4.**
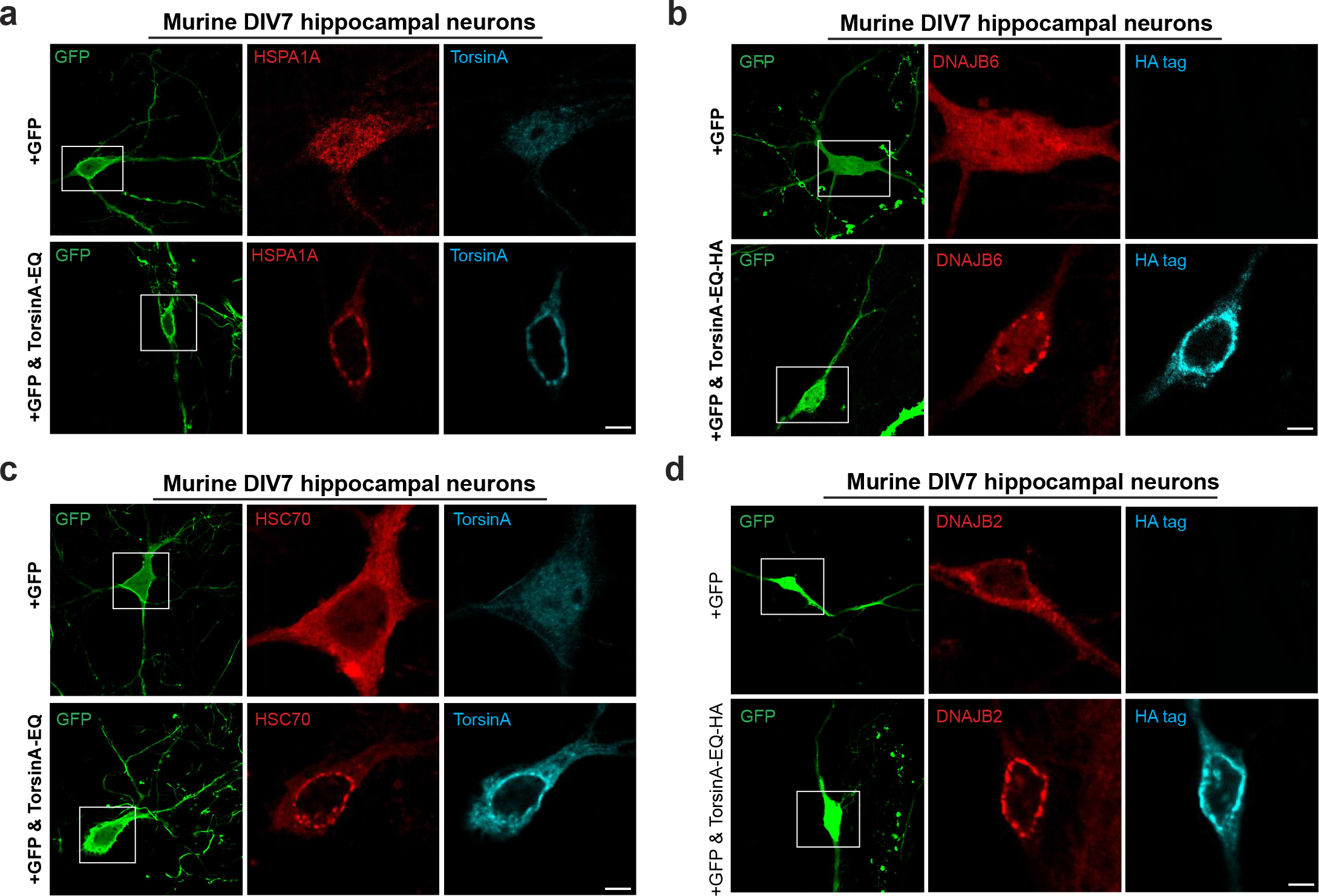
Highly abundant molecular chaperones are sequestered into NE blebs of primary mouse neurons with compromised TorsinA function. **a-d,** Murine DIV4 hippocampal neurons were transfected with GFP and empty vector (top row of all panels) or a dominant-negative TorsinA-EQ construct (bottom row of all panels). Constructs were allowed to express for 72 hours before processing the DIV7 cultures for IF. GFP expression was used to distinguish neurons from other cell types in the heterogeneous primary cell culture. Localization of the chaperones shown in panels (**a-d**) was probed using antibodies against the indicated endogenous chaperone (red). Untagged TorsinA-EQ was transfected in panels (a) and (**c**) and detected with a TorsinA antibody (cyan). TorsinA-EQ-HA was transfected in panels (**b**) and (**d**) and detected with an anti-HA antibody (cyan). Scale bar, 5 µm for all panels.

### MLF2 recruits DNAJB6 to blebs

To better understand a possible relationship between MLF2 and DNAJB6, we depleted MLF2 from TorsinKO cells (Fig. 5a). Upon MLF2 knockdown, K48- Ub and HSPA1A remained efficiently sequestered into NE foci but DNAJB6 was no longer recruited to blebs (Fig. 5a). Next, we performed a radioimmunoprecipitation of endogenous HSPA1A from metabolically ^35^S-labeled WT and TorsinKO cells under non-targeting (siNT) or siMLF2 conditions (Fig. 5b). Of the protein co-immunoprecipitating with HSPA1A, three detectable bands were unique or highly enriched in the TorsinKO siNT condition (Fig. 5b).

**Figure 5.**
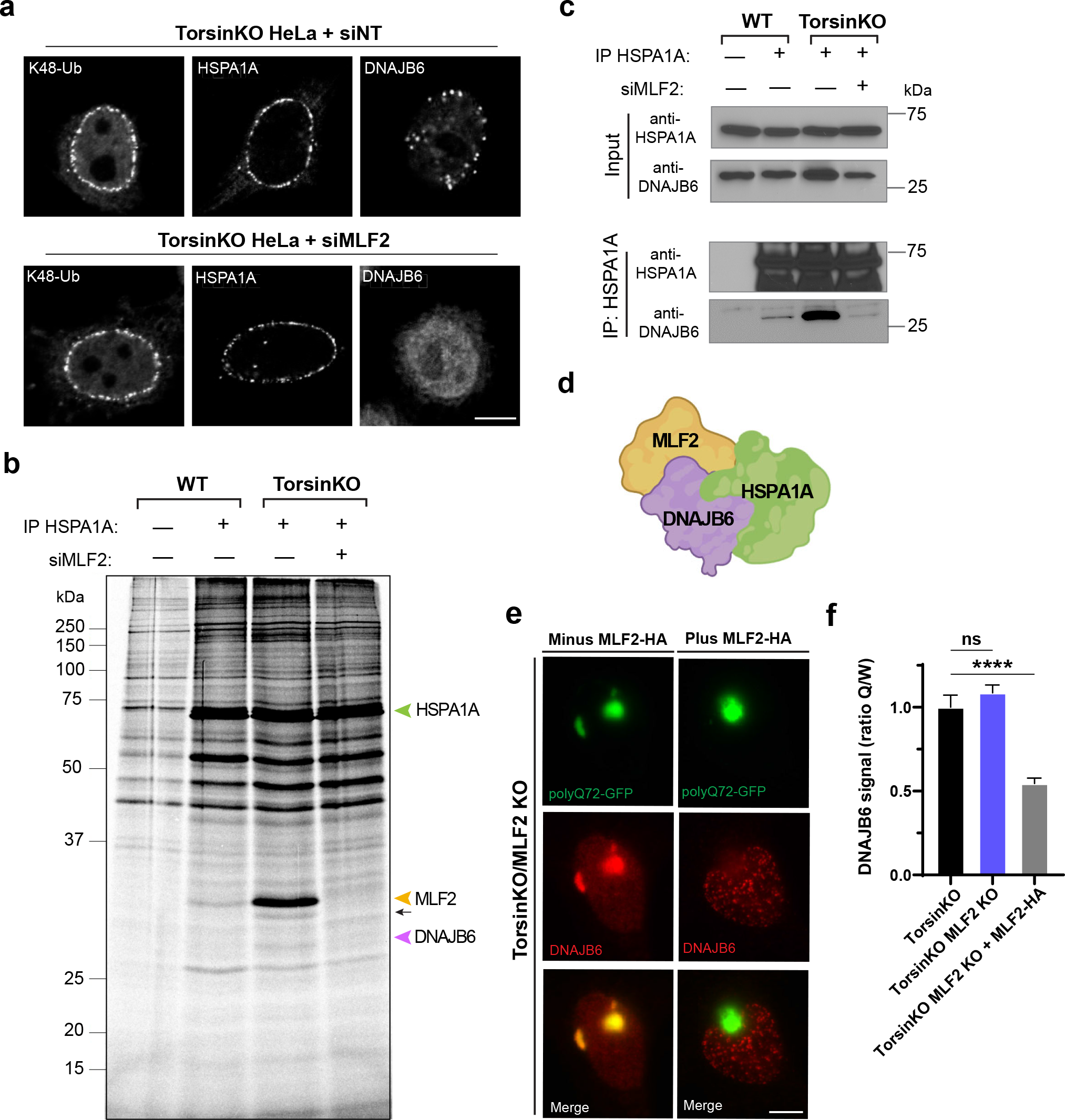
MLF2 is required for DNAJB6 to localize to NE blebs in Torsin-deficient cells. **a**, Representative IF images of chaperone localization upon knocking down MLF2. MLF2 depletion prevents DNAJB6 from localizing to blebs. Scale bar, 5 µm. **b**, WT and TorsinKO cells were metabolically labeled overnight with 150 µCi/mL 35S-Cys/Met and treated with a nontargeting or MLF2-targetting RNAi. HSPA1A was immunoprecipitated and stably associated proteins were detected by autoradiography. Green arrowhead, HSPA1A. Yellow arrowhead, MLF2. Black arrow, unknown protein. Purple arrowhead, DNAJB6. **c**, HSPA1A was immunoprecipitated from TorsinKO and WT HeLa cells under siNT or siMLF2 conditions. Note that HSPA1A retrieves far more DNAJB6 in TorsinKO cells compared to WT cells in an MLF2-dependent manner. **d**, Schematic model of the MLF2- DNAJB6-HSPA1A complex inside blebs. **e**, A “tug of war” experiment showing that overexpression of MLF2-HA in TorsinKO/MLF2KO cells titrates DNAJB6 out of polyQ72-GFP aggregates and into blebs. Representative IF images of TorsinKO/MLF2KO cells transfected with polyQ72-GFP alone (left column) or in combination with MLF2-HA (right column). Note that the HA channel is not shown. Scale bar, 5 µm. **f**, The ratio of DNAJB6 fluorescence signal inside polyQ72-GFP foci (Q) compared to whole cell (W) was calculated for 100 cells/condition. The data are shown as the mean ± standard deviation. Statistical analyses were performed using a two-tailed unpaired Student’s t-test where **** indicates p < 0.0001. ns, not significant.

Notably, HSPA1A interacted with more MLF2 in TorsinKO cells than in WT (Fig. 5b). Another band unique to the TorsinKO siNT condition migrates at the expected molecular mass of DNAJB6 and co-immunoprecipitates with HSPA1A exclusively in TorsinKO cells and in an MLF2-dependent manner (Fig. 5b). Additionally, a protein of unknown identity uniquely immunoprecipitates in an MLF2- and TorsinKO-dependent manner (Fig. 5b). Thus, HSPA1A interacts with DNAJB6 in an MLF2-dependent manner uniquely in TorsinKO cells.

To confirm that the MLF2-dependent co-immunoprecipitating protein was indeed DNAJB6, we performed a co-IP experiment (Fig. 5c). While a minor amount of DNAJB6 interacts with HSPA1A in WT cells, this interaction is significantly stabilized in TorsinKO cells in an MLF2-dependent manner (Fig. 5c). Based on these observations, we conclude that inside blebs, MLF2 and HSPA1A form a stable complex that interacts with DNAJB6 (Fig. 5d).

### The sequestration of chaperones in Torsin-deficient cells may contribute to proteotoxicity

DNAJB6 has a well-established role in preventing the formation of toxic inclusions, including polyglutamine (poly-Q) expansions that cause Huntington’s disease^38–40^. Thus, we were interested to know how this important role is affected by DNAJB6 sequestration into blebs. In TorsinKO/MLF2 KO cells, DNAJB6 is recruited to sites of poly-Q aggregation with high efficacy (Fig. 5e,f). However, upon re-introducing MLF2-HA via transient transfection, DNAJB6 is instead tightly sequestered into blebs (Fig. 5e,f). This redistribution of DNAJB6 away from a high affinity, aggregate-prone client underscores the pronounced proteotoxic potential of NE blebs with MLF2 being a critical modulator of this property.

### The sequestration of protein into NE blebs requires Nup98

While analyzing the APEX2- MLF2-HA MS datasets, we noticed that specific Nups were enriched in their proximity to MLF2 in TorsinKO cells including Nup50, Nup133, Nup98, and GP210 (Fig. 6a). While Nup50, Nup133, and GP210 integrate into stable subcomplexes of the NPC, Nup98 is found in multiple subcomplexes and binds the mRNA export factor Rae1^41^. Since the only protein known to localize to blebs in a K48-Ub-dependent manner is KSHV ORF10 (Fig. 1), which is known to form a complex with Nup98 and Rae1 ^32^ we prioritized our analysis on Nup98.

**Figure 6.**
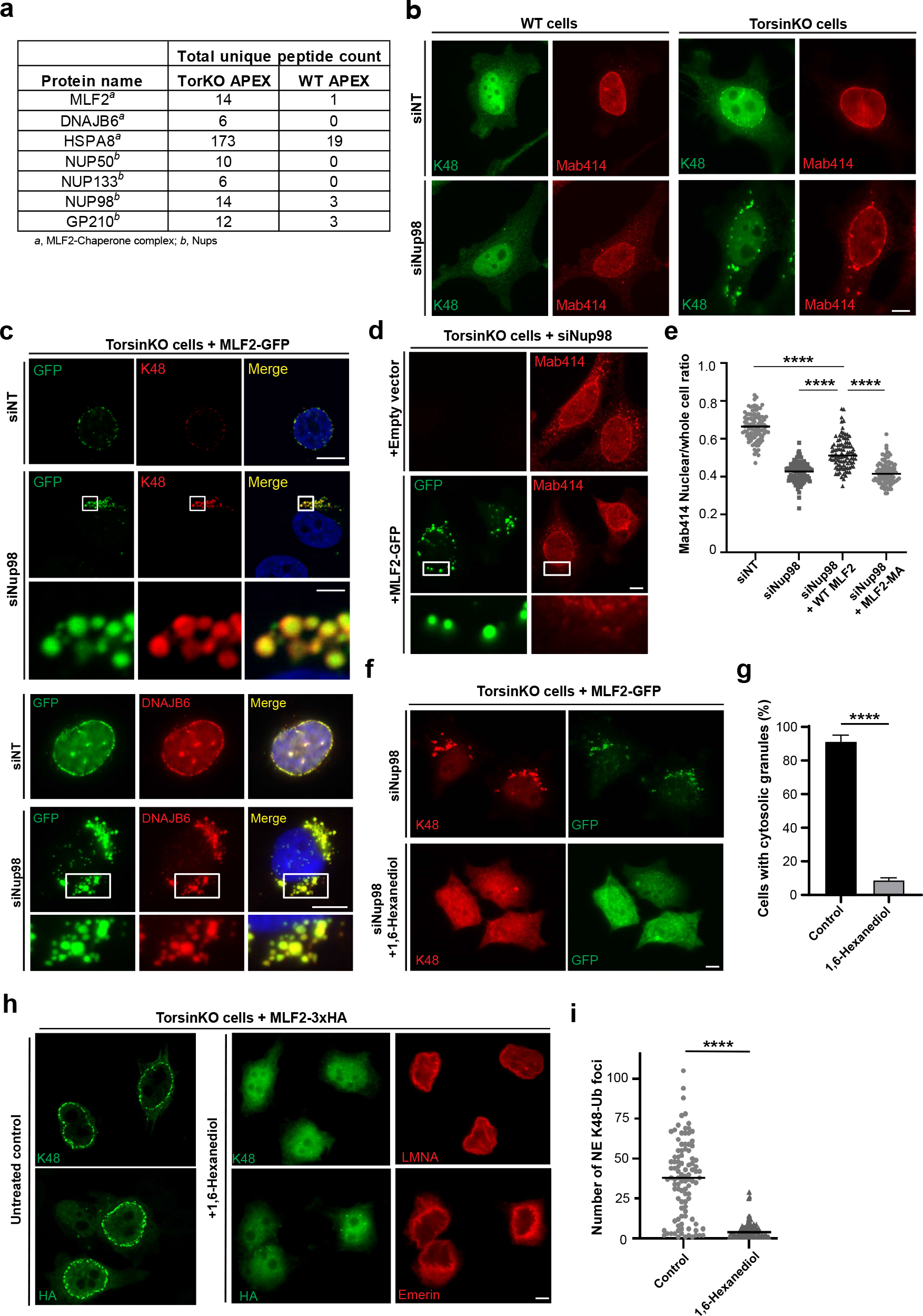
Nup98 is required for the NE sequestration of phase-separated granules composed of K48-Ub, FG-nucleoporins, MLF2, and chaperones. **a**, The proximity labeling MS strategy described in (Fig. 2a) reveals an enrichment of chaperones and nucleoporins interacting with MLF2 in TorsinKO cells. **b**, Representative IF images of WT and TorsinKO cells under siNup98 conditions. In TorsinKO cells, cytosolic granules enriched for K48-Ub and FG-Nups (Mab414, red) form upon siNup98. Note that the canonical NE accumulation of K48-Ub in TorsinKO cells is abolished under siNup98 conditions Scale bar, 5 µm. **c**, Representative IF images of TorsinKO cells expressing MLF2-GFP under siNup98 conditions. MLF2-GFP (top panel) localizes to the cytosolic granules containing K48-Ub that arise upon Nup98 depletion in TorsinKO cells. DNAJB6 (bottom panel) is also recruited to the cytosolic granules. Scale bar, 10 µm. **d**, Representative IF images of the effect on the FG-Nup accumulation in cytosolic granules upon overexpression of MLF2-GFP under siNup98 conditions. Scale bar, 5 µm. **e**, The ratio of nuclear to whole cell Mab414 signal was determined for 100 cells/condition. Expressing WT MLF2 significantly decreases the amount of cytosolic FG-Nup mislocalization upon siNup98. Statistical analyses were performed using a two-tailed unpaired Student’s t-test where **** indicates p < 0.0001. **f,** Representative IF images of TorsinKO cells expressing MLF2-GFP under siNup98 conditions in the absence (top row) or presence (bottom row) of 5% 1,6-hexanediol. Scale bar, 5 µm. **g**, The presence of cytosolic K48-Ub/MLF2-GFP granules upon Nup98 depletion was assessed for 100 cells/condition. The data are shown as the mean ± standard deviation. Statistical analyses were performed using a two-tailed unpaired Student’s t-test where **** indicates p < 0.0001. **h**, Representative IF images of TorsinKO cells expressing MLF2-3xHA (bottom row) in the absence (leftmost column) or presence (two rightmost columns) of 5% 1,6-hexanediol. NE integrity was monitored by laminA (LMNA, top) and emerin (bottom) staining. Scale bar, 5 µm. **i**, The number of K48-Ub foci around the NE rim was determined for 100/condition. Plots indicate the mean number of K48-Ub foci and statistical analyses were performed using a two-tailed unpaired Student’s t-test where **** indicates p < 0.0001.

Depleting Nup98 via siRNA provoked the formation of cytosolic granules composed of K48-Ub and FG-Nups to form exclusively in TorsinKO cells (Fig. 6b). We attribute this effect to Nup98 as this phenotype can be rescued by transfection of an siRNA-resistant Nup98 construct but not by Nup96, which is derived from a Nup98-96 precursor protein through proteolytic cleavage^42^ (Extended data 2a-d). We also examined whether other bleb components became incorporated into cytosolic granules upon Nup98 depletion. Both MLF2-GFP and DNAJB6 also localize to these cytosolic puncta (Fig. 6c) while HSPA1A and HSC70 did not (Extended data Fig. 2e). Together, we conclude that the FG-Nup Nup98 is required for the formation of an unusual NE compartment consisting of K48-Ub, MLF2, specific FG-Nups, and chaperones.

### Overexpressing MLF2 decreases the amount of mislocalized FG-Nups

When MLF2-GFP was overexpressed in TorsinKO cells in Nup98-depleted cells, we noticed a significant decrease in the amount of FG-Nup incorporation into the cytosolic granule (Fig. 6d). We calculated the nuclear/whole cell ratio of Mab414 signal in TorsinKO cells under siNT, siNup98, siNup98 + WT MLF2-FLAG, or siNup98 + MLF2^M/A^-FLAG (Fig. 6e). When Nup98 is depleted, significant FG-Nup mislocalization occurs and the nuclear/whole cell Mab414 ratio decreases (Fig. 6e). When WT-MLF2-FLAG is overexpressed in Nup98-silenced cells, the nuclear/whole cell Mab414 ratio significantly increases (Fig. 6e). This is not the case when MLF2^M/A^-FLAG is expressed (Fig. 6e), a mutant variant that fails to partition into blebs (see below, cf. Fig. 7b). Thus, MLF2 appears to exhibit a chaperone-like activity directed at cytosolic FG-Nups.

**Figure 7.**
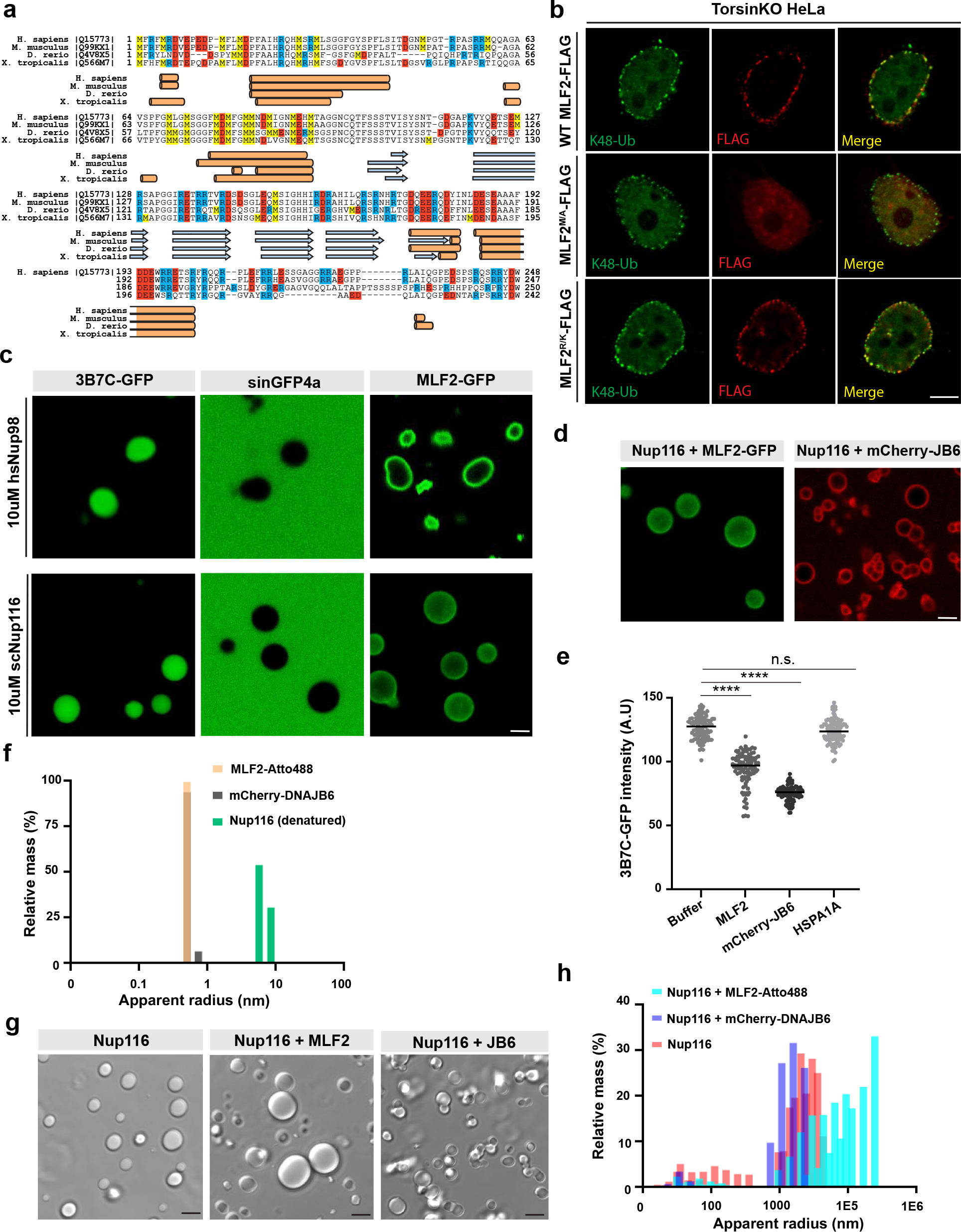
MLF2 localizes to phase separated condensates in vitro and modulates droplet permissiveness and size. **a**, Sequence alignment indicating that MLF2 is a methionine- and arginine-rich protein with a high degree of conservation. Methionine residues are highlighted in yellow boxes, arginine in blue, and positively charged residues in red. Orange cylinders indicate alpha helices predicted by Phyre261 and blue arrows represent predicted beta sheets. **b**, Representative IF images of MLF2-FLAG constructs in TorsinKO cells and their co-localization with K48-Ub. MLF2 requires methionine residues, but not arginine, to localize to blebs in TorsinKO cells. Scale bar, 5 µm. **c**, Purified MLF2-GFP interacts with phase separated droplets formed with Nup98 or Nup116 FG-domains. Purified, label-free Nup98 or Nup116 FG-domains were diluted from 300 µM stocks (in 2 M urea) to a final concentration of 10 µM in tris-buffered saline (TBS) to form droplets. Droplets were allowed to form at room temperature for five minutes before 3B7C-GFP, sinGFP4a, or MLF2-GFP was added to a final concentration of 5 µM. Scale bar, 2 µm. **d**, mCherry-DNAJB6 interacts with FG-domain droplets. 5 µM of MLF2-GFP or mCherry-DNAJB6 were introduced to 10 µM of Nup116 FG-domain droplets and imaged as described in panel (**c**). Scale bar, 2 µm. **e**, Nup116 droplets formed in the presence of untagged MLF2 or mCherry-DNAJB6, but not HSPA1A, are less permissible to 3B7C-GFP. 3B7C-GFP intensity was measured in the center of 100 droplets/condition. Plots indicate the mean intensity value and statistical analyses were performed using a two-tailed unpaired Student’s t-test where **** indicates p < 0.0001. **f**, Solutions of 5 µM purified MLF2-Atto488, mCherry-DNAJB6, or denatured Nup116 were analyzed by dynamic light scattering (DLS). Datasets composed of 100 reads (with five-second acquisition times) were collected for each condition. **g**, Representative phase contrast images of 5 µM Nup116 droplets formed in the presence of buffer alone, 5 µM MLF2-Atto488, or 5 µM mCherry-DNAJB6. Scale bar, 5 µm. **h**, Solutions of 5 µM Nup116 were formed in TBS without any additional protein (red bars), with 5 µM mCherry-DNAJB6 (dark blue bars), or 5 µM MLF2-GFP (light blue bars). Droplets were analyzed by DLS as described for panel (**f**).

### Blebs share properties with phase separated condensates

The cytosolic granules that form upon siNup98 in TorsinKO cells were often spherical (Fig. 6b-d), prompting us to ask whether they represent phase separated condensates. One strategy to probe the nature of such condensates is to determine their sensitivity to 1,6-hexanediol, which interrupts weak hydrophobic contacts and dissolves many phase separated condensates^43, 44^. When treated with 10% 1,6-hexanediol, the K48-Ub and MLF2-GFP granules typically observed in TorsinKO cells under siNup98 conditions largely dissolved (Fig. 6f,g).

The functional yeast ortholog of Nup98, *Saccharomyces cerevisiae* Nup116, serves as “Velcro” to recruit other FG-Nups during NPC assembly^45^. We therefore surmised that Nup98 similarly recruits FG-Nups to blebs, possibly to an abundance that allows for phase separation. Indeed, when we treat TorsinKO cells expressing MLF2-HA with 1,6-hexanediol, the K48-Ub and MLF2-HA are released from blebs despite the NE remaining intact (Fig. 6h,i). These observations are consistent with the interpretation that blebs represent phase separated entities composed of K48-Ub, MLF2, FG-Nups, and chaperones.

### MLF2 requires its high methionine content to localize to blebs

MLF2 possesses an unusually high arginine and methionine content, a property that is well conserved (Fig. 7a). While the average protein contains ∼4% arginine and <2% methionine, MLF2 is composed of 12.5% arginine and 7.7% methionine. Surface exposure of these two residues facilitates the diffusion of otherwise inert molecules into hydrogels^46^. Thus, it is possible that MLF2 utilizes its arginine and methionine content to immerse within the bleb phase.

We designed mutants of MLF2 that lack arginine (MLF2^R/K^) or methionine (MLF2^M/A^) and assessed their ability to localize to blebs (Fig. 7b). While the MLF2^R/K^ mutant readily co-localizes with K48-Ub foci around the nuclear rim, MLF2^M/A^ remains nucleoplasmic (Fig. 7b). These data suggest that MLF2 relies primarily on its methionine content to localize to blebs.

### MLF2 immerses into phase separated droplets *in vitro*

To investigate whether the recruitment of MLF2 relies on FG-driven phase separation, we purified FG domains from the *Homo sapiens* Nup98 and *the S. cerevisiae* Nup98 functional homolog Nup116. Under denaturing conditions (2M urea), Nup98 and Nup116 remain in a non-interacting state and do not phase separate^47^. However, upon dilution into buffer without denaturants these FG domains form cohesive interactions that produce phase separated droplets, which exhibit selective permeability^46, 47^ (Fig. 7c). We formed FG domain droplets and validated their selectivity with the purified GFP derivatives, 3B7C-GFP and sinGFP4a^46^ (Fig. 7c). 3B7C-GFP behaves like an NTR and readily partition into the FG-rich phase while sinGFP4a is designed to be “inert” and nearly completely exclude from the phase^46^ (Fig. 7c). Indeed, both Nup98 and Nup116 FG droplets allowed partitioning of 3B7C-GFP and exclusion of sinGFP4a (Fig. 7c).

We next purified MLF2-GFP and tested its ability to immerse into FG phases (Fig. 7c).

MLF2-GFP readily immersed into Nup116 droplets but remained mostly at the surface of Nup98 droplets (Fig. 7c). This difference in MLF2-GFP immersion is most likely due to the different organisms from which the FG domains are derived; Nup98 homologs from metazoans form very dense FG hydrogels that, in unmodified forms, are impermeable even to NTR-cargo complexes that readily pass through the NPC *in vivo*^48^. When Nup98 FG hydrogels are modified with O--N-acetylglucosamines (O-GlcNAc), as they are heavily *in vivo*, they allow rapid and full immersion of NTR-cargo complexes^48^. O-GlcNAc modifications are not found in lower eukaryotes such as yeast^49^. The FG domains used in this study were not glycosylated and therefore, the resulting Nup98 droplets likely prevent the full immersion of MLF2-GFP while the Nup116 droplets are more readily permeable by permissive cargo. Taken together, we conclude that MLF2 effectively partitions into FG-rich phases.

### MLF2 and DNAJB6 interact with FG phases and prevent the accumulation of an NTR-like GFP *in vitro*

We purified mCherry-DNAJB6 to address whether this chaperone also interacts with FG droplets. mCherry-DNAJB6 accumulates mainly on the surface of Nup116 droplets (Fig. 7d). An important consequence of MLF2 or mCherry-DNAJB6 interacting with Nup116 particles is that they prevent the full partition of 3B7C-GFP into the phase (Fig. 7e). This is not the case when droplets are formed in the presence of purified HSPA1A, which is excluded from the droplet (Fig. 7e, Extended data Fig. 3a). Thus, MLF2 and/or DNAJB6 impart a selectivity barrier to the phase separated droplets.

### MLF2 promotes the formation of large phase separated droplets *in vitro*

We noticed that droplets formed in the presence of MLF2 were significantly larger compared to other conditions (Fig. 7g). To quantify this effect, we performed dynamic light scattering (DLS) and examined the Nup116 particles’ apparent radii under different conditions. When solutions containing MLF2- Atto488, mCherry-DNAJB6, or denatured Nup116 alone were analyzed by DLS, small particles (<10 nm radius) corresponding to protein monomers were detected (Fig. 7f). Diluting scNup116 into non-denaturing buffer caused significantly larger particles (≥1,000 nm radius) to be detected as the phase separated droplets formed (Fig. 7h).

When Nup116 droplets were formed in the presence of mCherry-DNAJB6, we observed particles with a somewhat smaller size distribution compared to Nup116 alone (Fig. 7h).

However, when droplets were formed with MLF2-Atto488, a pronounced shift towards larger particle sizes occurred (Fig. 7h). This suggests that MLF2 facilitates the phase separating process, revealing a hitherto unknown FG-Nup directed activity.

## Discussion

In this study, we developed a virally-derived model substrate (Δ133 ORF10, Fig. 1) to explore the molecular composition and cellular consequences of nuclear envelope (NE) herniations that arise in disease models of primary dystonia. While NE blebs are ubiquitous in dystonia model organisms and are also found in a variety of other experimental and physiological settings^50^, it has generally been unclear whether NE blebs contribute to disease Δ133 ORF10 and the protein MLF2 in a comparative proteomics approach, we found that blebs are highly enriched for the FG-nucleoporin Nup98 and specific members of the HSP40 and HSP70 chaperone family including DNAJB2, DNAJB6, HSC70, and HSPA1A (Fig. 2). In cells with perturbed Torsin function, these chaperones become tightly sequestered within the lumen of NE blebs and are titrated away from their normal subcellular localization (Fig. 3). Importantly, this sequestration also occurs in primary murine neurons devoid of Torsin function (Fig. 4). Our finding that overexpressed chaperone constructs become efficiently sequestered into NE blebs (Extended data 1a,b) suggests that blebs have a remarkable capacity for chaperones. This raises the question of what mechanisms are at work to confer this unusual property.

We demonstrate that NE blebs contain FG-rich, phase-separated condensates (Fig. 6) that impose two proteotoxic challenges: an unprecedented degree of chaperone sequestration that is typically only observed upon overexpression of disease alleles, and a profound stabilization of normally short-lived proteins (Fig. 1e,f). We furthermore find that MLF2 promotes FG-domain condensation and enhances the specificity of clients partitioning into the phases (Fig. 5B, Fig. 7). Our observation that MLF2 expression results in a near-complete re- distribution of DNAJB6–a critical factor for suppressing proteotoxic aggregation^38, 40^– from poly-Q inclusions to NE blebs (Fig. 5e,f) establishes MLF2 as an important yet underappreciated player in protein homeostasis. Taken together, we uncover a direct proteotoxic contribution of blebs to DYT1 dystonia pathology and a function for MLF2 in altering the properties of phase separated NE compartments.

While nuclear transport receptors (NTRs) function as FG-Nup-directed chaperones during postmitotic NPC assembly^51^, interphase NPC biogenesis follows a distinct insertion pathway^52^. We demonstrate that MLF2 overexpression prevents the ectopic accumulation of FG-Nups upon Nup98 depletion (Fig. 6d,e). It is therefore tempting to speculate that non-NTR chaperones function during interphase NPC biogenesis to prevent non-productive interactions of FG-Nups destined for interphase assembly. As we demonstrate that MLF2 and DNAJB6 prevent the full partition of NTR-like molecules into FG-Nup phases (Fig. 7e), these chaperones may also prevent nuclear transport machinery from interacting prematurely with FG-Nups in nascent pores. As NE blebs result from stalled or defective interphase NPC biogenesis^23^, our observation that bulk nuclear transport cargo is excluded from FG-rich NE blebs^19, 23^ is consistent with this idea. It will be interesting for future studies to investigate whether and how the bleb-resident proteins identified herein (Fig. 2, Supplementary table 1) contribute to NPC assembly. Indeed, Kuiper et al. have independently assigned DNAJB6 to a role in NPC biogenesis (in press).

We have observed that MLF2-GFP accumulates at sites of NE membrane curvature in Torsin-deficient cells ^23^. This clustering of MLF2 juxtaposed against the curved NE may represent facilitated nucleation events of FG-Nups undergoing phase separation at sites of *de novo* interphase NPC assembly. This *in vivo* role for MLF2 would reflect the activity we report *in vitro* (Fig. 7g). Interestingly, MLF2 recruitment to blebs requires an unusually high methionine content (Fig. 7a,b). This sidechain can form bridging “aromate-methione-aromate” and other non-covalent interactions with aromatic residues^53^, which are abundant in FG-Nups. It will be interesting for future studies to investigate if other nuclear processes that rely on phase separation, or the phase-separating behavior of prion domain-containing proteins relying on aromatic “stickers”^54^, employ MLF2 activity and to explore whether these are compromised in diseases.

The observation that mutations in ER/NE-luminal Torsin ATPases give rise to an indirect proteotoxic mechanism across compartmental borders is unexpected. This feature adds a unique disease mechanism to the growing list of movement disorders with functional ties between liquid-liquid phase separation, nuclear transport machinery, and proteotoxicity^55–57^. Our finding that NE blebs exert a twofold proteotoxicity also represents a distinct pathological mechanism from the documented nuclear transport defects that arise in Torsin-deficient cells due to compromised NPC assembly^13, 21–23^ (Fig. 8).

**Figure 8.**
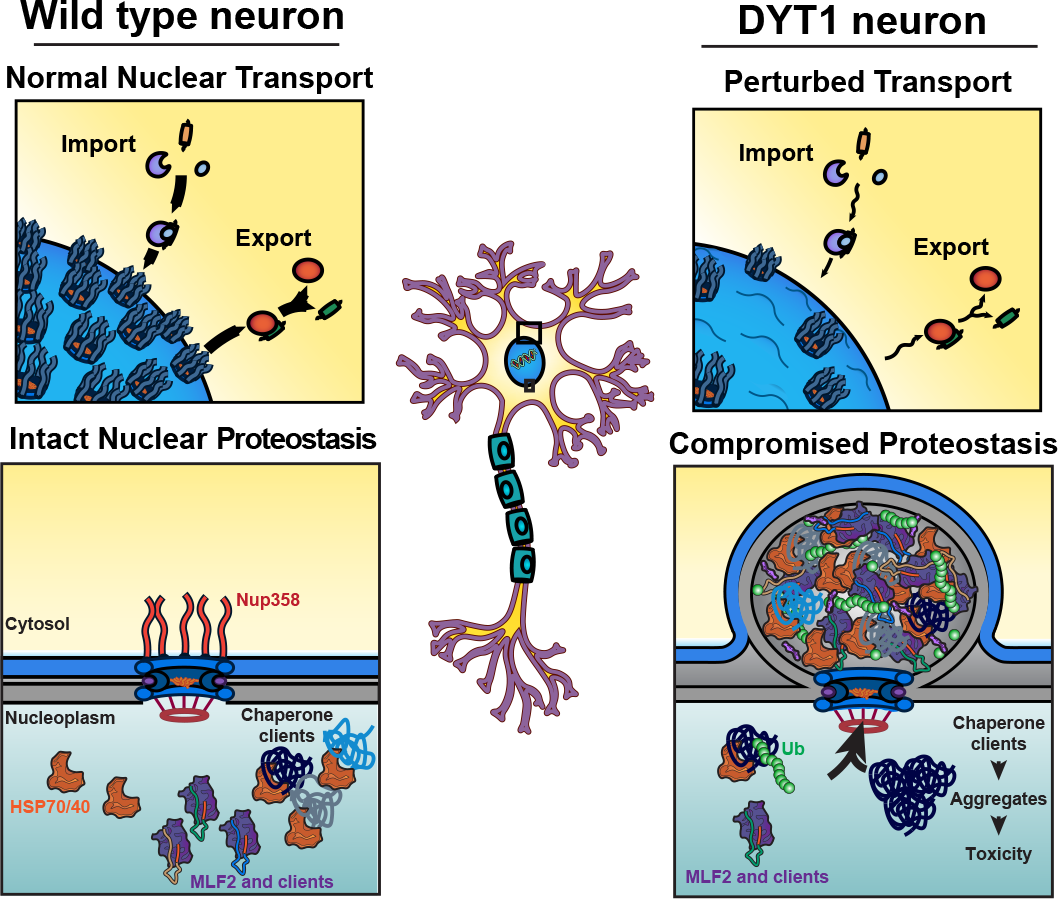
Dual proteotoxicity contributes to disease etiology in DYT1 Dystonia. Schematic model for how proteotoxicity may accumulate in Torsin- deficient cells. In wild type neurons, NPC biogenesis is unperturbed and chaperones are free to interact with clients. In DYT1 dystonia neurons, nuclear transport is perturbed due to defective NPC biogenesis. As FG-NUP containing blebs form instead of mature NPCs, they sequester proteins normally destined for degradation, chaperones, and MLF2 into phase separated granules. When essential chaperones are sequestered away from clients in Torsin-deficient cells, proteotoxic species may be allowed to form and persist to a greater extent than in cells with normal chaperone availability.

Finally, we propose that the dual proteotoxicity mechanism contributes to the unusual disease manifestation in DYT1 dystonia. Unlike most other congenital movement disorders, DYT1 dystonia is characterized by a childhood onset^58^ and incomplete disease penetration as only one third of all TorsinA mutation carriers develop the disease^12^. Adolescent carriers of the DYT1 mutation will never develop the disease if they have not developed symptoms prior to 30 years of age. These features may be explained by the fact that blebs are transient structures that resolve in juvenile model organisms^17^. Thus, the proteotoxicity imposed by blebs is likely confined to a specific window of neurological development. We propose that sequestered chaperones and accumulated short-lived proteins confer a window of vulnerability early in life that generates a high but potentially manageable degree of proteotoxic stress.

However, further insult on the PQC machinery that may normally be inconsequential could cause severe problems in these pre-sensitized cells. This model describes a stochastic and previously underappreciated influence on disease manifestation. While additional studies will be required to test these ideas in the future, our data provide a strong motivation to investigate pharmacological modulators of the cellular PQC system for DYT1 dystonia prevention and management.

## Online methods

### Antibodies and reagents

The following antibodies against the indicated proteins/epitopes were used in this study (WB, Western blot. IF, immunofluorescence): K48 linkage-specific polyubiquitin (WB, 1:4000. IF, 1:500. MilliporeSigma, Apu2), HA-peptide (WB, 1:2000. IF, 1:500. Roche, 3F10), p97 (WB,1:7000. Abcam, ab109240), α-tubulin (WB, 1:5000. MilliporeSigma, T5168), streptavidin-HRP (WB, 1:50000. Pierce, streptavidin Qdot 525 conjugate (IF, 1:500. Invitrogen, 10121013), HSPA1A (WB, 1:2000. IF, 1:500. Enzo, C92F3A-5), HSC70 (IF, 1:500. Santa Cruz, 7298), DNAJB6 (WB, 1:1000. IF, 1:500. Abcam, ab198995), DNAJB2 (IF, 1:500. Proteintech, 10838-1-AP), TorsinA (IF, 1:200. Homemade), Mab414 (IF, 1:500. Abcam, ab24609), GFP (IF, 1:500. Roche, 11814460001), emerin (IF, 1:1000. Santa Cruz, 25284), FLAG-peptide (IF, 1:500. MilliporeSigma, F3165), rabbit IgG HRP conjugate (WB, 1:10000. SouthernBiotech, 4030-05), mouse IgG HRP conjugate (WB, 1:20000. SouthernBiotech, 1030-05), rabbit & mouse IgG Alexa488 conjugates (IF, 1:700. Invitrogen, A11008 & A28175), rabbit & mouse IgG Alexa568 (IF, 1:700. Invitrogen, A-11011& A-11004), rabbit IgG Alexa633 (IF, 1:1000. Invitrogen, A- 21070).

Cycloheximide was purchased from Sigma-Aldrich (01810) and used at a final concentration of 100 µg/mL. Thymidine (Sigma-Aldrich, T1895) was used at a final concentration of 2.5 mM. Hexanediol was purchased from Millipore (804308) and used at 5% w/v.

### Plasmids, transient RNAi knockdowns, and transient transfections

KSHV ORF10-HA was custom synthesized by Genscript into the pcDNA3.1 vector. Δ ORF10-HA was cloned from the full lenth cDNA into pcDNA3.1. Plasmids containing the deubiquitylating enzyme M48^33^ were a gift from Hidde L. Ploegh (Whitehead Institute for Biomedical Research, Cambridge, MA). DNAJB2-HA, DNAJB12-HA, MLF2-3xHA and MLF2- GFP were cloned into pcDNA3.1 or pEGFP-N1 as previously described^23^. The plasmids containing His-tagged 3B7C-GFP, sinGFP4a, Nup98 and Nup116 FG domains were gifts from Dirk Görlich (Max Planck Institute for Biophysical Chemistry, Göttingen, Germany)^46, 47^.

To generate stable HeLa cell lines, MLF2-APEX2-HA was cloned into the pRetroX-Tight-Pur vector and 6 µg was transiently transfected into HEK293T cells along with 2 µg of MMLV gag/pol and 1 µg of the viral envelope protein VSV-G.

Nontargeting RNAi and RNAi targeting MLF2 and Nup98 were performed with SMARTpool oligos from Horizon Discovery. Knockdown efficiency was validated by quantitative PCR (qPCR) using iQ SYBR Green mix with a CFX Real-Time PCR 639 Detection System (Bio-Rad). For each knockdown, we employed the comparative Ct method using the internal control transcript RPL32. Primer sequences used for qPCR were as follows: MLF2 (FWD: GGACTC CCCTTCCCGACAGT, REV: GCCTCTCAGCCTGTACAAGAG)^23^, Nup98 (FWD: ACCACCCAGAACACTGGCTT, REV: GGCTGTGAGGCTTGGGTTAC). All primers were synthesized by Integrated DNA Technologies.

All plasmid transfections were performed with Lipfectamine 2000 (Invitrogen) according to manufacturer’s instructions and allowed to express for 24 hours prior to analyses. All RNAi knockdowns were performed with Lipfectamine RNAiMAX (Invitrogen) according to manufacturer’s instructions and allowed to incubate for 48 hours before analyses.

### Cell culture and cell lines

Torsin-deficient HeLa cells and their parental WT cells were cultured as previously described^19, 23^. Cells were maintained in Dulbecco’s modified Eagle’s medium (DMEM) supplemented with 10% v/v FBS (Thermo Fisher Scientific) and 100 U/mL of penicillin-streptomycin (Thermo Fisher Scientific). Cells were routinely checked for mycoplasma and determined to be free of contamination through the absence of extranuclear Hoechst 33342 (Life Technologies) staining.

Expi293 suspension cells were cultured in flat-bottom shaking flasks in preformulated Expi293 Expression Media (Gibco).

To generate stable HeLa cell lines expressing MLF2-APEX2-HA, retroviral vectors were produced in HEK293T cells via the transfection strategy described above. After 72 hours of expression, the supernatant was collected and filtered through a 0.45 µm filter before storage at -80°C. HeLa cells were seeded in 6-well plates and transduced with 100 µL viral supernatant plus 4 µg/mL polybrene reagent (Sigma-Aldrich). After 24 hours, media was switched to contain 1 µg/mL of puromycin (Sigma-Aldrich). Antibiotic selection was performed for 7 days before the dox-inducible MLF2-APEX2-HA expression was verified.

### CRISPR/Cas9 generation of MLF2 knockout

To generate MLF2 KO HeLa cells, we employed the CRISPR/Cas9 system^59^ as previously implemented^60^. Briefly, a guide RNA targeting MLF2 was cloned into the px459 vector (pSpCas9(BB)-2A-Puro (px459) V2.0 was a gift from Feng Zhang (Addgene plasmid #62988)). The sequence of this gRNA was as follows: sense 5’-CACCGGCTTCCATATCTTTCCAGTGA-3’ and antisense 5’-AAACCACTGGAAAGATATGGAAGCCA-3’. The MLF2 gRNA was cloned into the px459 vector and transfected into HeLa cells. The cells underwent antibiotic selection for 48 hours via treatment with 0.4 µg/mL puromycin (ThermoFisher). After selection, cells were seeded at a low density such that single-cell colonies could be isolated after 10 days in culture. These colonies were expanded and screened for MLF2 knockout by genotyping PCR as previously described^60^ with the following primers: FWD, CAACCATTCCTAGCAATGGG. REV, GGAAAGGACAGTGCTCTGAG.

### Hippocampal cultures, transfection, and immunofluorescence

Hippocampal cultures were performed on BALB/c P0 pups, in accordance with IACUC protocol number 2019-07912. Hypothermia was induced in pups, followed by decapitation and hippocampal dissection. Hippocampi were dissociated using 200 units Papain (Worthington LS003124, 200 units) and were plated on 14 mm Poly-D-lysine coated coverslips in a 24 well dish at a density of 150,000 cells per well. Neurons were transfected at DIV 4 with 1 µL per reaction of lipofectamine 2000 at a concentration of 0.6 µg per plasmid, for a total of 1.8 µg of DNA. Neurons were fixed at DIV 7 with 4% v/v paraformaldehyde (PFA) in PBS at room temperature for 15 minutes, permeabilized with 0.1% v/v Triton-X for 15 minutes, blocked with 1% w/v BSA for one hour, and stained with primary antibodies in blocking buffer overnight at 4°C (1:1000 rat HA, 1:200 rabbit TorsinA). Following the primary stain, coverslips were washed with PBST three times for 5 minutes, followed by a secondary antibody stain in blocking buffer for 1 hour at room temperature (1:1000 rat Alexa 568 and rabbit 633). Coverslips were washed with PBST three times for 5 minutes, stained with DAPI for 10 minutes at room temperature, washed with PBST three times for 5 minutes, and mounted in Aqua-Mount (Lerner Laboratories).

### Immunofluorescence and confocal microscopy

HeLa and SH-SY5Y cells were grown on coverslips and prepared for IF by fixing in 4% PFA (ThermoScientific) in phosphate-buffered saline (PBS) for 20 minutes at room temperature (RT). Cells were permeabilized in 0.1% Triton X-100 (Sigma-Aldrich) for 10 minutes at RT, then blocked in 4% bovine serum albumin (BSA) for 30 minutes. Primary antibodies were diluted into 4% BSA and incubated with coverslips for 45 minutes at RT. After extensive washing with PBS, fluorescent secondary antibodies were diluted in 4% BSA and incubated with coverslips for 45 minutes in the dark at RT. Cells were washed and stained with Hoechst 33342 (Life Technologies) before mounting onto slides with Fluoromount-G (Southern Biotech).

For hexanediol experiments, cells were incubated for five minutes with complete DMEM containing 5% or 10% w/v hexanediol (Millipore) before fixing in 4% v/v PFA and processing for IF as described above.

Phase separated droplets were imaged by spotting a 10 µL volume of the indicated conditions onto a glass bottom dish (WillCo Wells).

All images were collected on an LSM 880 laser scanning confocal microscope (Zeiss) with a C Plan-Apochromat 63×/1.40 oil DIC M27 objective using ZEN 2.1 software (Zeiss).

### Cycloheximide chase

WT and TorsinKO cells were plated in a 10 cm dish and transfected with Δ ORF10-HA. After 24 hours of expression, each cell line was trypsinized and split into two tubes. Tubes were incubated at 37°C with gentle shaking and treated with either DMSO or 100 µg/mL CHX and aliquots were taken at 0, 1, 2, 3, or 4 hours post treatment. Cells were collected via centrifugation and subjected to immunoblot for analyses.

### Immunoprecipitation, mass spectrometry preparation, and immunoblot analysis

For native IP experiments, cells were transfected with the indicated constructs 24 hours before harvesting. Cells were lysed in 1x NET buffer (150 mM NaCl, 50 mM Tris pH 7.4, 0.5% NP-40) supplemented with EDTA-free protease inhibitor cocktail (Roche) and 5 mM NEM (Sigma-Aldrich). Equal amounts of protein were loaded onto uncoupled protein G beads for one hour at 4°C to pre-clear lysates and reduce background contamination. Immunoprecipitation was conducted with pre-cleared lysates on anti-HA affinity matrix (Roche), magnetic beads conjugated to streptavidin, or magnetic protein G beads (Pierce) non-covalently coupled to anti-HSPA1A. Protein was allowed to complex with the beads for three hours at 4°C before extensive washing with NET buffer. Stable interactions were eluted for immunoblot analyses in 30 µL of 2x SDS reducing buffer and heated at 70°C for five minutes. For downstream mass spectrometry applications, protein complexes were briefly run into SDS-PAGE gels, stained with SimplyBlue Safe Stain (ThermoFisher) before bands of 2-4 mm were extracted. Gel bands were submitted to the MS & Proteomics Resource at the Yale University Keck Biotechnology Laboratory where the samples were further processed for liquid chromatography-MS/MS analysis.

Radioimmunoprecipitations were carried out in as described previously^4, 61^. Briefly, metabolically labeled cells were grown in complete DMEM containing 150 µCi/mL ^35^S-Cys/Met labeling mix (PerkinElmer) for 16 hours prior to lysis in NET buffer. Stably associating complexes were immunoprecipitated as described above. Co-eluting protein were detected by autoradiography and imaged on a Typhoon laser-scanning platform (Cytiva).

Immunoblotting was carried out with IP eluates or cell lysates in supplemented NET buffer. Equal micrograms of protein were resolved in SDS-PAGE gels (Bio-Rad) and transferred onto PVDF membranes (Bio-Rad). Membranes were blocked in 5% w/v milk in PBS + 0.1% Tween- 20 (Sigma-Aldrich). Primary and HRP-conjugated secondary antibodies were diluted in blocking buffer. Blots were visualized by chemiluminescence on a ChemiDoc Gel Imaging System (Bio-Rad).

### APEX2 reaction and NE enrichment

To induce the expression of MLF2-APEX2-HA, cells were treated with 500 µg/mL dox for 24 hours. After 24 hours, cells were incubated for 30 minutes with complete DMEM containing 500 µM biotin phenol. To conduct the APEX2 reaction, cells were treated with 1 mM H_2_O_2_ for one minute before the reaction was quenched by washing the plates three times with quenching buffer (PBS, pH 7.4, 0.5 mM MgCl_2_, 1 mM CaCl_2_, 5 mM Trolox, 10 mM sodium ascorbate, 10 mM sodium azide). Cells were collected via trypsinization and enriched for NE fractions as described previously^23, 62^. Briefly, cells were gently pelleted in buffer containing 250 mM sucrose and homogenized via trituration through a 25-guage needle. The homogenates were layered onto a 0.9 M sucrose buffer and spun down. The pellets (membrane fractions and nuclei) were solubilized overnight in buffer without sucrose containing benzonase, heparin, NEM, and protease inhibitors. The solubilized nuclei were spun at 15,000 x g for 45 minutes and the supernatant was collected as the nucleoplasm and pellet as the NE/ER enriched fraction. The ER/NE fraction was solubilized in 8 M urea and equal amounts of protein were loaded onto streptavidin beads for capture of biotinylated protein, which were analyzed by immunoblot or mass spectrometry as described above.

### Transmission electron microscopy and immunogold labeling

Electron microscopy (EM) and immunogold labeling was performed as previously described^19, 23^ at the Yale School of Medicine’s Center for Cellular and Molecular Imaging. Briefly, cells were fixed by high-pressure freezing (Leica EM HPM100) and freeze substitution (Leica AFS) was carried out at 2,000 pounds/square inch in 0.1% uranyl acetate/acetone. Samples were infiltrated into Lowicryl HM20 resin (Electron Microscopy Science) and sectioned onto Formvar/carbon-coated nickel grids for immunolabeling.

Samples were blocked in 1% fish skin gelatin, then incubated with primary antibodies diluted 1:50 in blocking buffer. 10 nm protein A-gold particles (Utrecht Medical Center) were used to detect the primary antibodies and grids were stained with 2% uranyl acetate and lead citrate.

Image were captured with an FEI Tecnai Biotwin TEM at 80Kv equipped with a Morada CCD and iTEM (Olympus) software.

### Recombinant protein expression and purification

Nup98 and Nup116^46, 47^ were purified from BL21(DE3) *E. coli* strains under denaturing conditions by virtue of an N-terminal His_18_ tag. Transformed BL21(DE3) cells were induced with mM isopropyl- -D-thiogalactoside (IPTG) at an optical density (OD) of 0.6 and allowed to express for 16 ^β^s at 16°C in terrific broth media (Sigma-Aldrich). Cell pellets containing His-Nup116 were resuspended in room temperature native lysis buffer (50 mM Tris, pH 7.5, 300 mM NaCl, 10 mM β Me, 2 uL benzonase, 1 mM PMSF) while His-Nup98 pellets were resuspended in denaturing lysis buffer (8 M urea, 50 mM Tris, pH 8, 20 mM imidazole, 10 mM β-me, 1 mM PMSF). Resuspended cell pellets passaged twice through a French Press.

Following a 15-minute centrifugation at 20,000 x g, His-Nup116 was solubilized from inclusion bodies by resuspending the pellet in denaturing lysis buffer. His-tagged Nups were complexed with Ni-NTA agarose (Qiagen) at room temperature for 1 hour before the columns were extensively washed with wash buffer (6 M urea, 50 mM Tris, pH 8, 25 mM imidazole, 10 mM β me). Protein was eluted in elution buffer (6 M urea, 50 mM Tris, pH 8, 400 mM imidazole, 10 mM β me) and dialyzed overnight at 4°C into UTS buffer (2 M urea, 50 mM Tris, pH 7.4, 150 mM NaCl). Finally, the dialyzed Nup was concentrated using an Amicon Ultra-4 10,000 MWCO centrifugation unit (MilliporeSigma).

Phase separated droplets were formed by diluting 300 µM Nup stock solutions in denaturing UTS buffer into tris-buffered saline(TBS; 50 mM Tris, pH 7.4, 150 mM NaCl) to a final concentration of 5 or 10 µM. Droplets spontaneously form upon rapid dilution out of urea^47^.

mCherry-DNAJB6-His_10_ was purified as previously described^63^. BL21(DE3) *E. coli* transformed with Cherry-DNAJB6 were induced with 0.5 mM IPTG at an OD of 0.6 and allowed to express for 16 hours at 16°C in terrific broth media (Sigma-Aldrich). Cell pellets were resuspended in ice cold resuspension buffer (100 mM Tris pH 8, 150 mM NaCl, 10 mM β-Me, 1 mM PMSF) and were passaged through the French Press twice. Lysates were clarified by a 15-minute 20,000 x g spin and mCherry-DNAJB6-His_10_ was solubilized from inclusion bodies in pellet buffer (100 mM Tris pH 8, 8 M urea, 150 mM NaCl, 20 mM imidazole, 10 mM β-Me, 1 mM PMSF). mCherry-DNAJB6-His_10_ was allowed to bind Ni-NTA agarose via batch mode purification for 1 hour at 4°C. The matrix was washed extensively in wash buffer (100 mM Tris pH 8, 8 M urea, 150 mM NaCl, 20 mM imidazole, 10 mM β-Me) and mCherry-DNAJB6-His_10_ was eluted in elution buffer (100 mM Tris pH 8, 150 mM NaCl, 10 mM concentration with an Amicon Ultra-4 10,000 MWCO centrifugation unit (MilliporeSigma). The protein was buffer-exchanged into final buffer (50 mM Tris pH 7.5, 150 mM KCl).

His-tagged GFP variants were purified as previously described^46^. Briefly, BL21(DE3) cells transformed with His-brSUMO-sinGFP4a or His-brSUMO-3B7C-GFP were induced with 0.5 mM IPTG at an OD of 0.6 and allowed to express at 37°C for three hours in terrific broth (Sigma-Aldrich). Cell pellets were resuspended in resuspension buffer (50 mM Tris pH 7.5, 150 mM NaCl, 10 mM β Me, 20 mM imidazole, 1 mM PMSF) and passaged through the French Press twice. Lysates were clarified by spinning for 15 minutes at 20,000 x g. His-tagged proteins were bound to Ni-NTA agarose (Qiagen) for 1 hour at 4°C before washing in resuspension buffer.

GFP constructs were eluted in resuspension buffer containing 400 mM imidazole, then concentrated with an Amicon Ultra-4 10,000 MWCO centrifugation unit (MiliporeSigma). Final concentrates were buffer-exchanged into TBS.

HSPA1A-His_10_-FLAG was purified from BL21(DE3) cells induced with 0.5 mM IPTG at an OD of and allowed to express for 16 hours at 16°C in terrific broth media (Sigma-Aldrich). Pellets were resuspended in resuspension buffer (100 mM HEPES pH 8, 500 mM NaCl, 10% glycerol, 10 mM β-Me, 20 mM imidazole, 1 mM PMSF) and passed through the French Press twice. After clarifyi ^β^ the lysate, the supernatant was applied to Ni-NTA agarose (Qiangen) for 1 houra t 4C.

The matrix was washed extensively in wash buffer (30 mM HEPES pH 7.4, 500 mM NaCl, 10% v/v glycerol, 20 mM imidazole, 10 mM β-Me) and protein was eluted in wash buffer with 400 mM imidazole. Protein was concentrated ^β^ buffer-exchanged as described above in to TBS.

MLF2 constructs (MLF2-GFP-FLAG and MLF2-Atto488) were purified from a mammalian Expi293 suspension system. cDNA was cloned into a pcDNA3.1 vector and transfected into Expi293 cells using the ExpiFectamine 293 Transfection Kit (Gibco). Cells were harvested after 72 hours of expression and pellets were lysed in lysis buffer (50 mM Tris pH 7.5, 150 mM NaCl, 1% DDM, 10% glycerol, protease inhibitor tablet (Roche)). After clarifying the lysate, the supernatant was applied to anti-FLAG M2 Affinity Gel (Sigma-Aldrich) and allowed to bind overnight at 4°C. The matrix was washed extensively with wash buffer (50 mM Tris pH 7.5, 2 mM ATP, 150 mM NaCl, 0.05% DDM) then washed with wash buffer minus detergent. Protein was eluted by incubating the matrix with final buffer (50 mM Tris pH 7.5 150 mM NaCl, 0.3 mg/mL FLAG peptide) for 1 hour at 4°C. MLF2 constructs were concentrated as described above and buffer-exchanged into TBS. All purified proteins were aliquoted and flash frozen in liquid nitrogen before long-term storage at -80°C.

### Atto-tagging via the sortase reaction

To produce MLF2- and HSPA1A-Atto488, constructs were purified with an LPETG sortase recognition sequence between the C-terminal end and the downstream purification tag. This motif is recognized by the transpeptidase Sortase, which catalyzes a reaction wherein a molecule harboring a poly-Glycine label is attached to the LPETG sequence^64^. Reactions were carried out using purified MLF2-LPETG-FLAG or HSPA1A-LPETG-His and the Sortag-IT™ ATTO 488 Labeling Kit (Active Motif) according to manufacturer’s instructions. After the sortase reaction, free dye was removed from the Atto-tagged proteins by washing extensively with TBS in Amicon Ultra-0.5 mL Centrifugal Filters (MilliporeSigma), then running through a PD MiniTrap desalting column (Cytiva).

### Dynamic light scattering

DLS was used to assess the distribution of FG particles forming the in presence or absence of MLF2. Measurements were taken on a DynaPro Titan DLS instrument (Wyatt Technologies) at 25°C and data were analyzed using DYNAMICS software (Wyatt Technologies). 10 µL reactions of 5 µM Nup116 in TBS containing no additional protein, MLF2-Atto488, or mCherry- 797 DNAJB6 were allowed to form for five minutes before diluting to 50 µL volume. Note that Nup116 was added to TBS containing the additional proteins, i.e., Nup116 droplets were formed 799 in the presence of MLF2 or DNAJB6 constructs. The 50 µL volume was transferred to a quartz cuvette and datasets of 100 measurements of five-second acquisition times were collected.

### Image processing and statistical analyses

All images were analyzed with Fiji software^65^. To quantify the percent of TorsinKO cells with NE foci of various protein (Fig. 5a,f), the nucleus was selected as the region of interest (ROI) and the number of foci was quantified in Fiji as previously performed^19, 23^ using the “Find Maxima” function with a noise tolerance of 10. Cells with ≥ foci were considered to have foci while those with <20 were not as occasional foci within the nucleoplasm occur even in WT cells. The foci status of 100 cells/condition was determined. Whether cells had cytosolic granules under siNup98 conditions (Fig. 6g) was determined by visualizing the presence or absence of cytosolic K48-Ub deposits for 100 cells/condition.

To quantify the rescue effect of MLF2 on FG-Nup mislocalization (Fig. 6e), we imaged 100 cells/condition and quantified the Mab414 antibody signal intensity in the whole cell and the nucleus by selecting these as ROIs in Fiji. The ratio of nuclear Mab414 intensity to whole cell intensity was calculated.

3B7C-GFP intensity was calculated inside Nup116 droplets by selecting the center of droplets as an ROI, then quantifying the average GFP signal intensity in Fiji. 100 droplets were quantified for each condition.

All measurements were taken from distinct samples or cells. Statistical analyses were performed in GraphPad Prism. For normally distributed datasets, which we confirmed using a Shapiro-Wilk test, statistical significance was assessed via unpaired two-tailed parametric t tests. Data that were not normally distributed were tested for statistical significance using a non-parametric two-tailed Mann Whitney test.

## Acknowledgements

This work is supported by NIH R01GM114401 (C.S.), DOD PR200788 (C.S.), NIH 5T32GM007223-44 (S.M.P.), NIH F31NS120528 (S.M.P.), NIH R56MH122449 (A.J.K.), NIH R01MH115939 (A.J.K.), NIH NS105640 (A.J.K.), NIH F31MH116571 (J.E.S.) and the Dystonia Medical Research Foundation (C.S. and A.J.R.). We thank Dirk Gorlich and members of his laboratory for sharing reagents. We also thank the Yale Keck Biophysical Resource Center Mass Spectrometry & Proteomics Resource as well as the Biophysical Resource Laboratory. We thank Morven Graham and the Yale Center for Cellular and Molecular Imaging for assistance with electron microscopy.

## Author contributions

S.M. Prophet, A.J. Rampello, and C. Schlieker conceptualized and designed experiments in the text. S.M. Prophet, A.J. Rampello, J.E. Shaw, and R.F. Niescier performed experiments. S.M. Prophet, A.J. Rampello, J.E. Shaw, R.F. Niescier, A.J. Koleske, and C. Schlieker analyzed and interpreted data. S.M. Prophet, and C. Schlieker wrote the original manuscript. S.M. Prophet, A.J. Rampello, J.E. Shaw, R.F. Niescier, A.J. Koleske, and C. Schlieker revised and edited the manuscript.

---

**Extended data 1.**
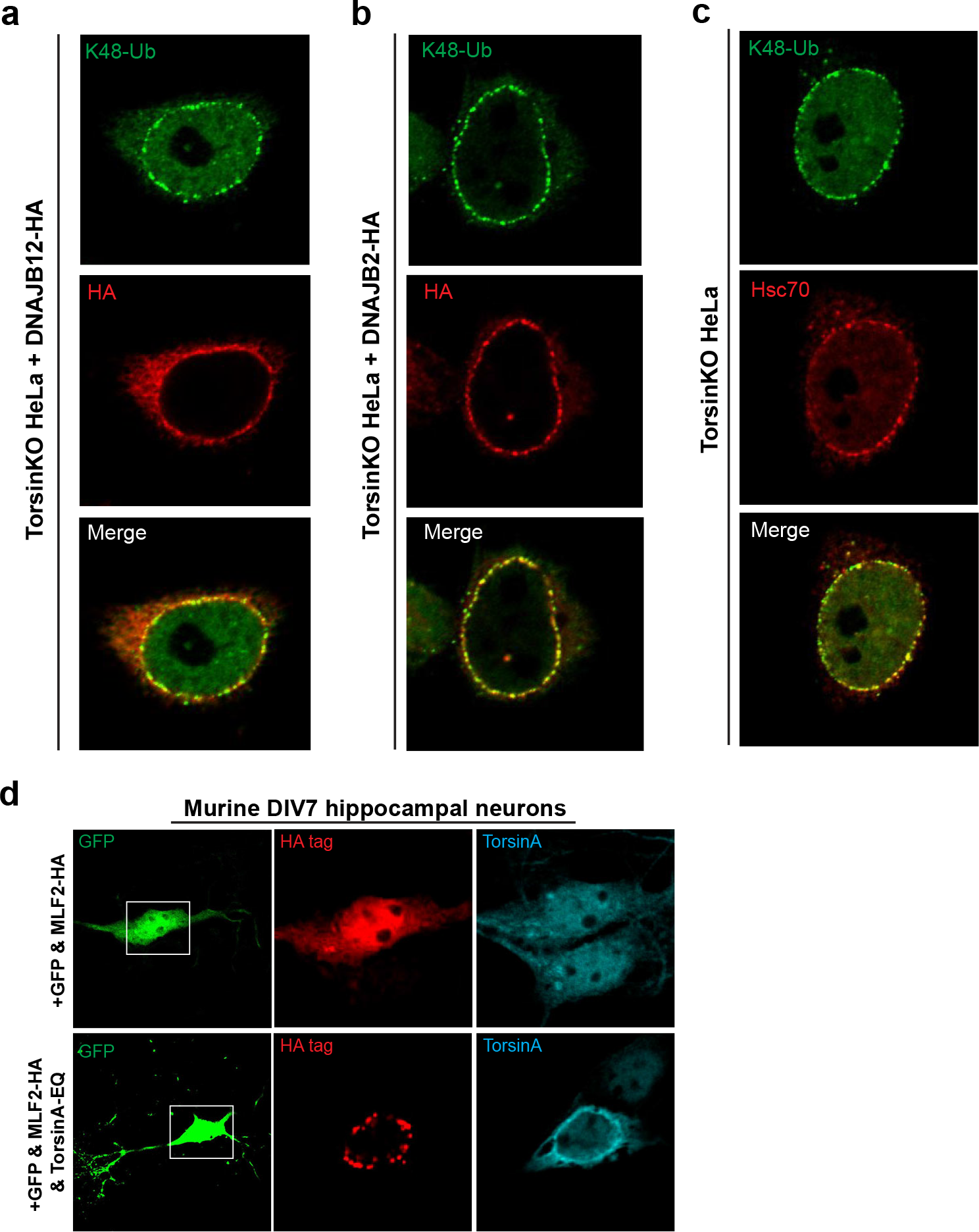
Validation of candidates identified by the MS datasets by IF. **a**, A representative IF image of overexpressed DNAJB12-HA in TorsinKO cells. DNAJB12-HA has a weak propensity to co-localize with K48-Ub foci at the NE. **b**, A representative IF image of overexpressed DNAJB2-HA in TorsinKO cells. As we have previously observed (ref.23), DNAJB2-HA has a strong propensity to co-localize with K48-Ub foci at the nuclear rim. **c**, Representative image of endogenous HSC70 co-localizing with K48-Ub foci in TorsinKO cells at the nuclear rim. **d**, Murine DIV4 hippocampal neurons were transfected with GFP and either MLF2-HA alone or in combination with a dominant-negative TorsinA-EQ construct. Constructs were allowed to express for 72 hours before processing the DIV7 cultures for IF. Note that GFP expression was used to distinguish neurons from other cell types in the heterogeneous primary cell culture.

**Extended data 2.**
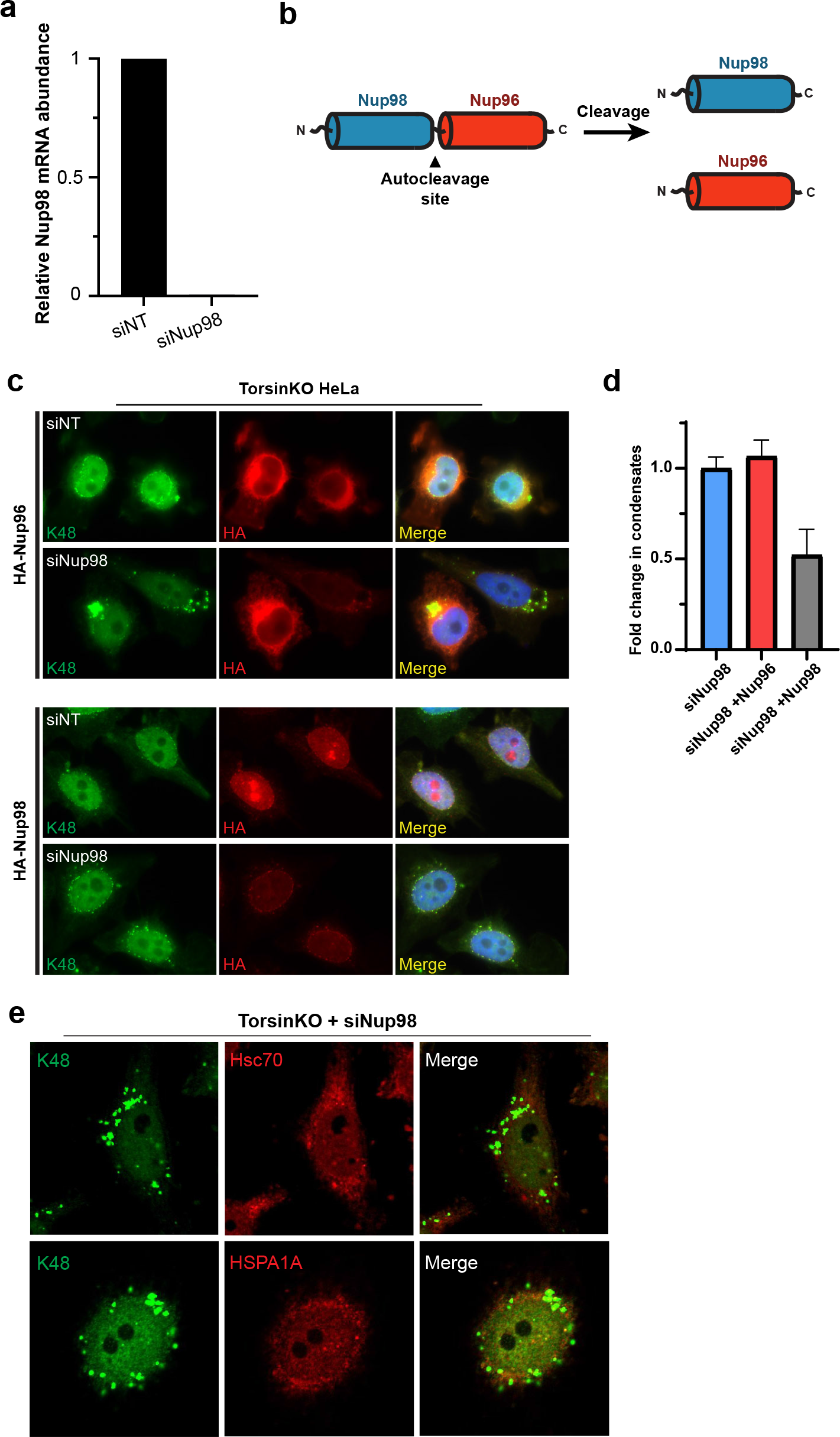
Validation of Nup98 knockdown and HPC70 localization. **a**, qPCR validation of Nup98 depletion upon 48 hours of 50 nM siRNA treatment. Relative Nup98 transcript levels are normalized to RPL32. **b**, Nup98 and Nup96 are translated as a single precursor protein that undergoes an autocleavage event to produce the two individual proteins (ref.42). Thus, RNAi knockdown of Nup98 results in the simultaneous depletion of Nup96. **c**, Representative IF images of TorsinKO cells expressing HA-Nup96 or HA-Nup98 under nontargeting or siNup98-96 conditions. To distinguish which protein’s knockdown produces the cytosolic granules in TorsinKO cells, HA-tagged Nup98 or Nup96 was assessed for the ability to rescue the phenotype under knockdown conditions. Note that the K48-Ub cytosolic granules are not produced under siNup98 when HA-Nup98 is expressed. **d**, Quantification of the rescue affect when HA-Nup96 or HA-Nup98 are expressed. The presence of cytosolic inclusions was assessed for 100 cells/ condition and normalized to the untransfected knockdown control. The data are shown as the mean ± standard deviation. **e**, Representative IF images of Hsc70 and HSPA1A localization upon Nup98 depletion.

**Extended data 3.**
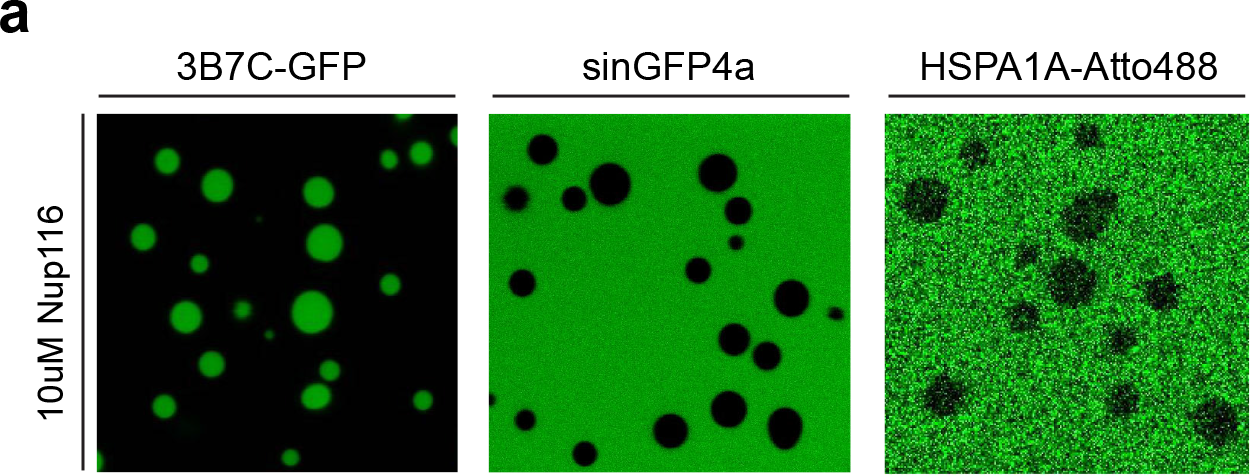
HSPA1A is excluded from Nup116 droplets. **a**, Purified HSPA1A was tagged with an Atto488 label and incubated with Nup116 droplets. The selective permeability of the droplets was confirmed by the NTR-like molecule 3B7C-GFP and the excluded GFP variant, sinGFP4a.

